# Predicting agronomic traits and associated genomic regions in diverse rice landraces using marker stability

**DOI:** 10.1101/805002

**Authors:** Oghenejokpeme I. Orhobor, Nickolai N. Alexandrov, Dmytro Chebotarov, Tobias Kretzschmar, Kenneth L. McNally, Millicent D. Sanciangco, Ross D. King

## Abstract

To secure the world’s food supply it is essential that we improve our knowledge of the genetic underpinnings of complex agronomic traits. In this paper, we report our findings from performing trait prediction and association mapping using marker stability in diverse rice landraces. We used the least absolute shrinkage and selection operator as our marker selection algorithm, and considered twelve real agronomic traits and a hundred simulated traits using a population with approximately a hundred thousand markers. For trait prediction, we considered several statistical/machine learning methods. We found that some of the methods considered performed best when preselected markers using marker stability were used. However, our results also show that one might need to make a trade-off between model size and performance for some learning methods. For association mapping, we compared marker stability to the genome-wide efficient mixed-model analysis (GEMMA), and for the simulated traits, we found that marker stability significantly outperforms GEMMA. For the real traits, marker stability successfully identifies multiple associated markers, which often entail those selected by GEMMA. Further analysis of the markers selected for the real traits using marker stability showed that they are located in known quantitative trait loci (QTL) using the QTL Annotation Rice Online database. Furthermore, co-functional network prediction of the selected markers using RiceNet v2 also showed association to known controlling genes. We argue that a wide adoption of the marker stability approach for the prediction of agronomic traits and association mapping could improve global rice breeding efforts.

## Background

Rice accounts for approximately 20% of global dietary energy needs, feeding more than half of the world’s population [14], making it the most important cereal crop. The world’s population is predicted to increase by almost three billion by 2050 [27], implying that we will need to double rice yield [33]. However, breeding for rice yield potential is stagnant, as the highest yielding rice varieties were introduced about 30 years ago [11]. Furthermore, the challenge of increasing rice yield is complicated by climate change, which will increase biotic and abiotic stresses [35]. We contend that the best hope of overcoming breeding stagnation is to be found in recent technological advances of genomic selection (GS), allele mining in large diverse populations, discovering new gene-trait associations and causative variations, and using genome editing for introducing desirable alleles into elite lines.

### Predicting agronomic traits

Complex phenotypes can be successfully predicted using statistical/machine learning methods. The incorporation of these predictions in accelerated breeding methods is known as GS; as breeders make their selection of progenitor based on the prediction results rather than physically observed phenotypes. In GS genomic estimated breeding values (GEBVs) are assigned to individuals using models that estimate the relationship between a population’s genetics and phenotypes of interest. GS requires a dense marker coverage of the genome, with all markers being considered simultaneously in the model learning process. This differentiates GS from traditional marker assisted selection (MAS), which is limited to using just a few predefined markers [20]. GS therefore relies on modern sequencing and genotyping technologies that enable the provision of hundreds of thousands genetic markers in populations of interest.

One consequence of dense marker coverage of the genome is that the number of markers (*p*) is usually much larger than the number of individuals (*n*), (*p* ≫ *n*). This makes it more difficult to identify markers that are significantly linked to an agronomic trait within a given population. Genome wide association studies (GWAS) aim to discover key markers associated with phenotypes. These studies are usually performed in populations with unrelated germplasm to maximize allele diversity [12, 18, 52, 49]. They also require that the genome of individuals in a population be densely covered by genetic markers, typically single nucleotide polymorphisms (SNPs). Within rice the use of GWAS has identified large effect genetic regions or quantitative trait loci (QTLs) associated with agronomic traits such as grain yield, plant height, flowering time, aluminium tolerance and submergence tolerance [4, 9, 12, 25, 46].

### Predicting agronomic traits through associated genomic regions in rice

In this study we investigate the use of marker stability for the identification of associated markers in rice. To do this we used the diverse population of the 3,000 rice genomes project [3], using twelve rice agronomic traits, and a hundred simulated traits. In a standard GWAS procedure [19, 29, 30], markers are tested in isolation and a multiple testing correction procedure is then used to control for false positives [5, 6].

In contrast, we propose the use of marker stability for the identification of associated markers. Marker stability in this context is equivalent to feature stability in the machine learning literature [28]. In machine learning, marker selection is used to identify markers that are strongly associated with a trait using a marker selection algorithm. This procedure is typically done using all samples in a population simultaneously. However, the markers that are selected in this case may not be stable. That is, should there be a change to the individuals in the population, the set of markers that are selected by a marker selection algorithm might be slightly different. Therefore, the goal of marker stability is to identify a set of markers that are consistently associated with a trait irrespective of changes to the individuals in a population.

Stable markers are typically identified by subsampling a given population and performing marker selection using a marker selection algorithm. The markers that are most frequently deemed important in each subsampled set are then considered stable in the work done in [26] on stability selection. An extension to the work in [26] was reported in [2]. The authors demonstrate that the markers selected by stability selection may be too conservative and may miss regions known to be associated with a trait, although it effectively controls the family-wise error rate. They argue that a group of SNPs, rather than single SNPs should be considered when performing association mapping by selection stability. Furthermore, ensemble methods which identify stable markers through a consensus of parametric and non-parametric algorithms has also been proposed [1]. We adopt the approach by [26] in this study using the least absolute shrinkage and selection operator [40, 41] (LASSO) as our marker selection algorithm.

Here, we performed our subsampling using *k*-fold cross-validation, and focussed on two marker sets. The first is the set of all uniquely identified associated markers selected in each cross-validation subsampled set, we refer to these markers as the aggregate cross-validation markers (ACVM). The second is the set of all markers that are deemed important in every cross-validation subsampled set, we refer to these markers as the intersect cross-validation markers (ICVM). We argue that the ICVM set does not contain markers that are associated by chance for two reasons. The first being that each marker in the marker set is selected because of its relationship to the others, and the second being that they are high frequency markers, consistently identified as associated with a trait of interest in every subsampled set. A known limitation of the use of LASSO is that only markers with linear additive effects are selected, thus ignoring non-linear epistatic interactions. However, we argue that the ICVMs represent a core set of associated markers, and by proxy, associated genomic regions that are present in all varieties in a diverse population. This is invaluable, as it lays the foundation for further investigation in sub-populations with exclusive epistatic interactions.

In addition to LASSO, we performed trait prediction using all available markers, the ACVMs and ICVMs with ridge regression (RR), ridge regression best linear unbiased predictor [15] (RBLUP), random forests [7] (RF) and gradient boosted machines [13] (GBM). We found that for some of the traits considered, all methods except RBLUP performed best when only preselected, associated markers are used. To check the biological relevance for the discovered markers we cross-referenced the intersect markers with known QTLs in the QTL Annotation Rice Online (Q-TARO) database [50]. For the twelve agronomic traits considered, and at different linkage disequilibrium (LD) thresholds, we found that many of the markers are in LD with known controlling genes. Furthermore, we statistically demonstrated that these associations are non-random. This strongly suggests that these markers not in LD with a known controlling gene are in previously unknown associated regions. Finally we compared marker stability to genome-wide efficient mixed-model analysis [53] (GEMMA), our results show that marker stability significantly outperforms GEMMA on the real and simulated traits.

## Methods

### Genotype and phenotype data

We used the Core SNP subset of 3000 Rice Genomes data version 0.4, comprising 996,009 SNPs from the International Rice Research Institute (IRRI). It was downloaded from http://snp-seek.irri.org and contains filtered SNP set (fraction of missing data <20%, minor allele frequency >1%, LD pruned with *r*^2^ of 0.8). An LD-pruned dataset with 101,595 markers was created using Plink [31] with a window of 50 SNPs, a step size of 5, and *r*^2^ value of 0.02. We converted each SNP call for all varieties to numeric values; class 1 homozygotes are represented with 1, class 2 homozygotes as -1, and heterozygotes with 0. Missing values were imputed using column means, as the effect of imputation in this case is minimal but is required by some of the prediction techniques considered.

Twelve real traits were considered with varying numbers of varieties due to the degree of missing values for some (Table 3). The trait data is taken from the public morpho-agronomic data on progenitor accessions in trials conducted by the International Rice Genebank at IRRI as part of the routine characterization of genetic resources. Most of the traits were expected to be highly heritable since they are standard morphological/agronomic descriptors used by genebanks for routine characterization. Hence, though data are from different years of screening without replication, most values will not show significant variation due to environmental differences. We did not impute missing values for phenotypes to avoid biasing the predictions. See Figure 4 for the distribution of the traits.

We simulated a hundred traits by randomly selecting a hundred SNPs from the genotype data also used for the real traits. For the SNPs selected for these traits, a weight *w* is assigned, where *w* is a value between 0 and 1, such that it follows a pareto distribution with a few SNPs having large effects. Trait values were computed for the simulated traits using the genotype data *g* for the randomly selected SNPs, such that

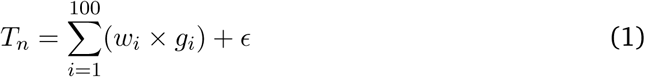

where *ϵ* ∈ 𝒩(3, 0.023). All 3023 varieties present in the original genotype dataset was used.

### Predictive models

The LASSO, proposed by Tibshirani, is a learner that reduces the number of markers when applied to GS by assigning some marker coefficients to zero. This is achieved using an *l*_1_ penalty which estimates LASSO coefficients, 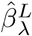, that minimize

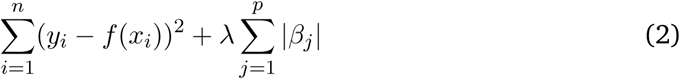

where *f*(*x*_*i*_) and *y*_*i*_ are the predicted and actual values for the *i*th variety in the population, referred to as the residual. The squared sum of all residuals for all varieties in the population is the residual sum of squares. *λ*∑_*j*_ |*β*_*j*_| is the shrinkage penalty and *λ* is the regularization parameter, which controls the impact of the shrinkage penalty on the regression coefficients. RR is similar to the LASSO with the difference being that it uses an *l*_2_ penalty. For both these methods, the regularization parameter was chosen using internal 10-fold cross validation.

RBLUP does not require any parameters to be tuned. For RF, the default of 1/3 the total number of variables is considered at each split, five observations are used for each terminal node, and 752 trees were grown for each forest when predictive models were built. When RF is used to rank selected SNPs 1000 trees are grown. For GBM a shrinkage parameter of 0.1 is used due to size of the datasets, an interaction depth of 6, and 1501 trees were grown. All predictive experiments were performed using R [39] and model performance was estimated as the coefficient of determination (*R*^2^).

### Experimental setup

For experiments using real traits we split the dataset into training and testing sets, 65% and 35% respectively with random sampling. A training/test data split was preferred over cross validation due to the high computational cost of the experiments. For LASSO-directed genomic predictions, we performed 3-fold cross validation on the training set to identify the associated markers. Two marker subsets using the non-zero markers from each fold were then created. The first is an aggregate of all uniquely selected markers from each fold, the ACVM set, and the second is an intersection of all markers from each fold, the ICVM set.

## Results

### Genomic selection using marker stability

We identified the associated ACVM and ICVM sets for twelve real agronomic traits, then performed genomic predictions with the standard dataset with 100k markers, ACVMs and ICVMs using the learning methods outlined in the previous section. We compared learner performance on the ACVM dataset to those when all 100k markers are used. The null hypothesis was that there is no difference in learner performance between the datasets (significance level of 0.05). A sign tests indicates that the null hypothesis can be rejected for RR and RBLUP, but not for LASSO, RF, and GBM with p-values of 0.006, 0, 0.388, 1, and 0.146 respectively. We observed that RBLUP’s performance reduced on the ACVM dataset for all traits, while RF’s improved for most. This suggests that reducing the number of markers eliminates noise (markers with no signal), which aids RF – but by doing so, small effect markers that aid RBLUP were also eliminated. For most traits, the ACVM’s performance is either marginally worse-off or outperforms the 100k dataset depending on the learner (Fig. 1). The slight performance loss may be a trade-off worth making as the models are easier to understand given the number of markers used (Table 1).

**Table 1:** Number of markers selected for each LASSO cross validation selection mathod. ACVM - Aggregate cross validation markers. ICVM - Intersect cross validation markers. These numbers are in comparison to the 101,595 markers in the complete marker set.

| Trait | ACVM | ICVM |
| --- | --- | --- |
| Culm diameter | 80 | 2 |
| Culm length | 147 | 24 |
| Culm number | 75 | 3 |
| Grain length | 109 | 10 |
| Grain width | 296 | 27 |
| Grain weight | 153 | 16 |
| Heading date | 197 | 25 |
| Ligule length | 191 | 4 |
| Leaf length | 137 | 13 |
| Leaf width | 212 | 13 |
| Panicle length | 158 | 15 |
| Seedling height | 44 | 1 |

**Table 2:** P-values of obtaining the same genes in LD with the selected markers (ICVM) by randomly sampled markers over 10,000 iterations at multiple LD thresholds. ICVM - Intersect cross validation markers. “-” is given in cases were no genes are in LD with the ICVMs.

| Trait | 100 Kbps | 500 Kbps | 1 Mbps | 1.5 Mbps | 2 Mbps |
| --- | --- | --- | --- | --- | --- |
| Culm diameter | 0.001 | 0.005 | 0.007 | 0.013 | 0.019 |
| Culm length | 0 | 0 | 0 | 0 | 0 |
| Culm number | - | 0.007 | 0.009 | 0.012 | 0 |
| Grain length | 0.002 | 0 | 0 | 0 | 0 |
| Grain width | 0.006 | 0 | 0 | 0 | 0 |
| Grain weight | 0 | 0 | 0 | 0 | 0 |
| Heading date | - | 0 | 0 | 0 | 0 |
| Ligule length | 0.001 | 0 | 0 | 0 | 0 |
| Leaf length | 0.005 | 0.034 | 0.048 | 0 | 0 |
| Leaf width | 0 | 0 | 0 | 0 | 0 |
| Panicle length | 0.004 | 0 | 0 | 0 | 0 |
| Seeding height | - | 0.002 | 0.004 | 0.005 | 0.006 |

**Table 3:** The number of varieties for the considered agronomic traits.

| **Trait** | **Definition** | **No of varieties** |
| --- | --- | --- |
| Culm diameter | Culm diameter (mm) of basal internode at re-productive | 1899 |
| Culm length | Culm length (cm) at reproductive | 1900 |
| Culm number | Culm number (count) at reproductive cultivated | 1901 |
| Grain length | Grain length (mm) | 2261 |
| Grain width | Grain width (mm) | 2261 |
| Grain weight | 100-grain weight (gm) cultivated | 2259 |
| Heading date | Heading (days to 80% fully headed) cultivated | 2265 |
| Ligule length | Ligule length (mm) cultivated | 1878 |
| Leaf length | Leaf length (cm) cultivated | 1901 |
| Leaf width | Leaf width (cm) cultivated | 1901 |
| Panicle length | Panicle length (cm) at post-harvest | 1896 |
| Seedling height | Seedling height (cm) | 1877 |

**Figure 1:**
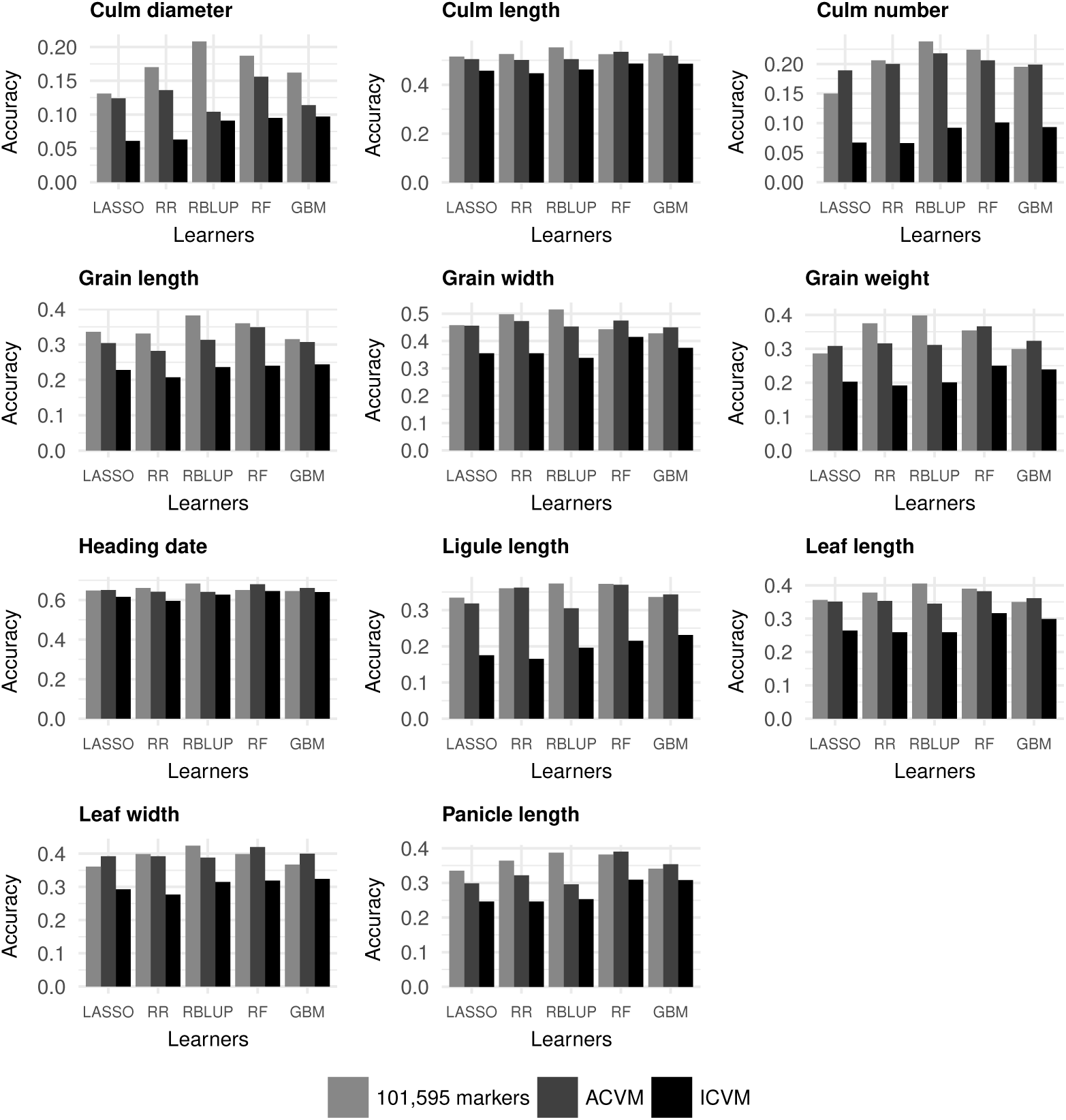
Predictive accuracies (*R*^2^) for the five learners considered. Predictive accuracy for LASSO, RR, RBLUP, RF, and GBM for all twelve traits using all markers and LASSO-Selected markers (ICVM and ACVM). ICVM - Intersect cross validation markers. ACVM - Aggregate cross validation markers. Seedling height is excluded because only one marker was selected in ICVM. Its accuracy using simple linear regression is 0.062.

We observed that traits for which a relatively small number of associated markers are identified have the lowest average correlation between observed and predicted values, even on the 100k dataset: for example, culm diameter and seedling height (Tables 1 and 4). The selection of few markers suggests that the trait is simple and controlled by only a few major QTLs, which suggest that these traits should be easy to predict. However this is not the case, and we have two hypotheses to explain this: that interactions that are unique to certain subpopulations might have been lost due to model generalization, and alternatively that these traits are mostly controlled by QTLs with non-linear relationships – we believe to be more probable.

**Table 4:**
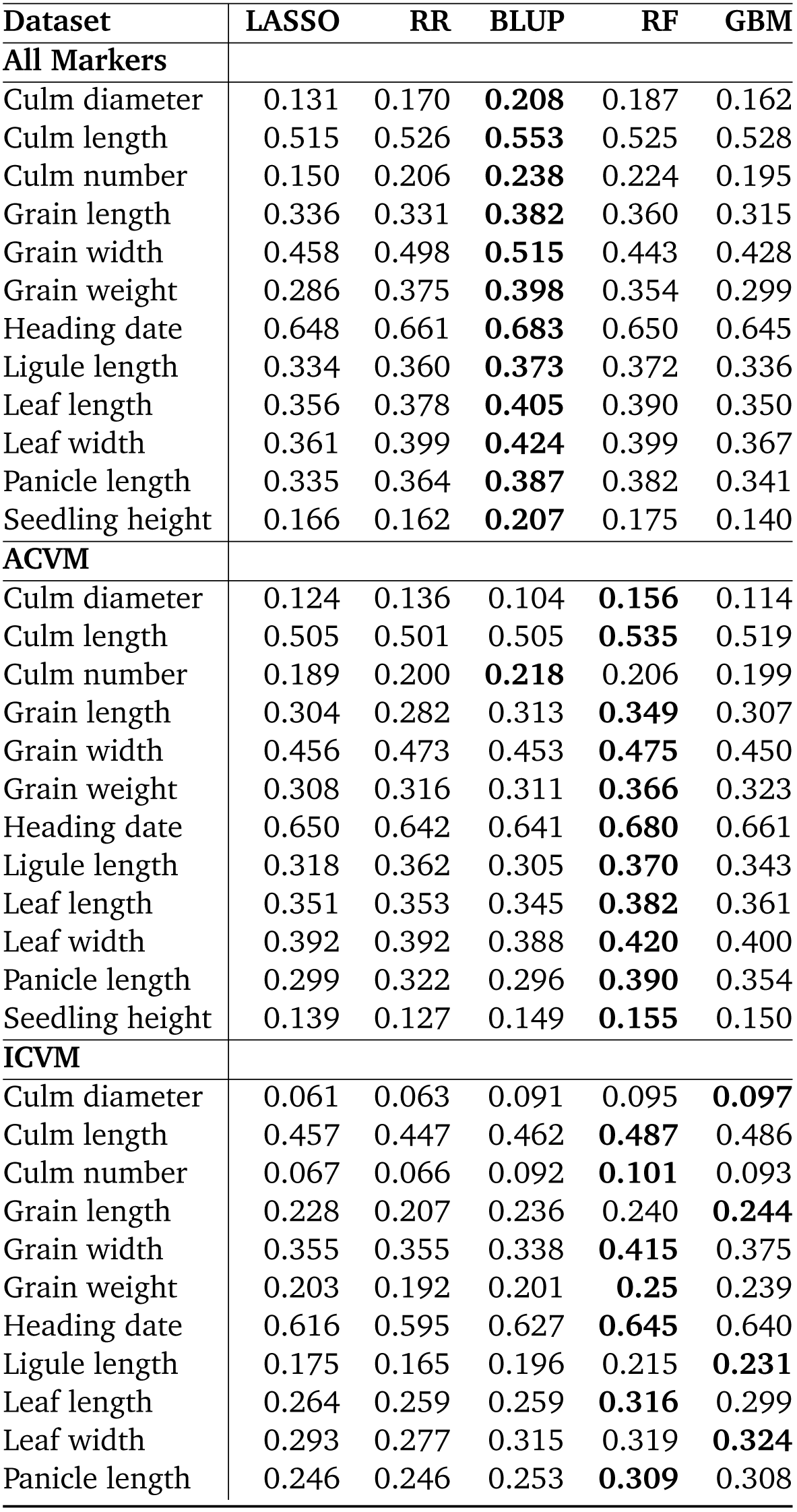
Predictive accuracy (*R*^2^) for LASSO, ridge regression, BLUP, random forests, and gradient boosted machines for all twelve real traits on the dataset with 105, markers and the LASSO-Selected markers (ACVM and ICVM). ACVM - Aggregate cross-validation markers. ICVM - Intersect cross-validation markers.

| Dataset | LASSO | RR | BLUP | RF | GBM |
| --- | --- | --- | --- | --- | --- |
| **All Markers** |  |  |  |  |  |
| Culm diameter | 0.131 | 0.170 | **0.208** | 0.187 | 0.162 |
| Culm length | 0.515 | 0.526 | **0.553** | 0.525 | 0.528 |
| Culm number | 0.150 | 0.206 | **0.238** | 0.224 | 0.195 |
| Grain length | 0.336 | 0.331 | **0.382** | 0.360 | 0.315 |
| Grain width | 0.458 | 0.498 | **0.515** | 0.443 | 0.428 |
| Grain weight | 0.286 | 0.375 | **0.398** | 0.354 | 0.299 |
| Heading date | 0.648 | 0.661 | **0.683** | 0.650 | 0.645 |
| Ligule length | 0.334 | 0.360 | **0.373** | 0.372 | 0.336 |
| Leaf length | 0.356 | 0.378 | **0.405** | 0.390 | 0.350 |
| Leaf width | 0.361 | 0.399 | **0.424** | 0.399 | 0.367 |
| Panicle length | 0.335 | 0.364 | **0.387** | 0.382 | 0.341 |
| Seedling height | 0.166 | 0.162 | **0.207** | 0.175 | 0.140 |
| **ACVM** |  |  |  |  |  |
| Culm diameter | 0.124 | 0.136 | 0.104 | **0.156** | 0.114 |
| Culm length | 0.505 | 0.501 | 0.505 | **0.535** | 0.519 |
| Culm number | 0.189 | 0.200 | **0.218** | 0.206 | 0.199 |
| Grain length | 0.304 | 0.282 | 0.313 | **0.349** | 0.307 |
| Grain width | 0.456 | 0.473 | 0.453 | **0.475** | 0.450 |
| Grain weight | 0.308 | 0.316 | 0.311 | **0.366** | 0.323 |
| Heading date | 0.650 | 0.642 | 0.641 | **0.680** | 0.661 |
| Ligule length | 0.318 | 0.362 | 0.305 | **0.370** | 0.343 |
| Leaf length | 0.351 | 0.353 | 0.345 | **0.382** | 0.361 |
| Leaf width | 0.392 | 0.392 | 0.388 | **0.420** | 0.400 |
| Panicle length | 0.299 | 0.322 | 0.296 | **0.390** | 0.354 |
| Seedling height | 0.139 | 0.127 | 0.149 | **0.155** | 0.150 |
| **ICVM** |  |  |  |  |  |
| Culm diameter | 0.061 | 0.063 | 0.091 | 0.095 | **0.097** |
| Culm length | 0.457 | 0.447 | 0.462 | **0.487** | 0.486 |
| Culm number | 0.067 | 0.066 | 0.092 | **0.101** | 0.093 |
| Grain length | 0.228 | 0.207 | 0.236 | 0.240 | **0.244** |
| Grain width | 0.355 | 0.355 | 0.338 | **0.415** | 0.375 |
| Grain weight | 0.203 | 0.192 | 0.201 | **0.25** | 0.239 |
| Heading date | 0.616 | 0.595 | 0.627 | **0.645** | 0.640 |
| Ligule length | 0.175 | 0.165 | 0.196 | 0.215 | **0.231** |
| Leaf length | 0.264 | 0.259 | 0.259 | **0.316** | 0.299 |
| Leaf width | 0.293 | 0.277 | 0.315 | 0.319 | **0.324** |
| Panicle length | 0.246 | 0.246 | 0.253 | **0.309** | 0.308 |

Depending on the trait, we observed that the ICVM dataset’s performance is between 25% to 95% of the ACVM’s. This is because the aggregate markers are more representative of the genetic diversity in the population. However, the intersect markers are significant irrespective of subpopulation, suggesting that these markers which can be interpreted as QTLs are significantly linked to the traits considered, making them predictors of associated QTLs.

### Identifying associated regions

We cross-referenced the positions of ICVMs identified for the real traits with QTLs in the Q-TARO database to validate their selection by determining if they are in LD with known controlling QTLs. We searched for associated QTLs with major and minor characteristics in Table 5. We considered multiple LD distance thresholds as there is no consensus in the literature as to how far a causal gene can be from a significant marker [8]. Five distance thresholds were considered; 100 Kbps, 500 Kbps, 1 Mbps, 1.5 Mbps, and 2 Mbps. At the most conservative threshold of 100 Kbps, some of the selected markers are in LD with a known controlling gene for six of the twelve traits, and at 1 Mbps, all traits have a selected marker in LD with a known controlling gene. It is worth noting that at 2 Mbps, just one of the twelve traits have selected markers that are all in LD with a gene of interest (Table 6). This suggests that the markers that are not in LD with known controlling genes are either in previously unknown associated regions, or that they perturb the markers that are in LD with known controlling genes.

**Table 5:** Trait characteristics used in marker validation.

| **Trait** | **Major character** | **Minor character** |
| --- | --- | --- |
| Culm diameter | Morphological trait | Culm leaf |
| Culm length | Morphological trait | Culm leaf |
| Culm number | Morphological trait | Culm leaf |
| Grain length | Morphological trait | Seed |
| Grain width | Morphological trait | Seed |
| Grain weight | Morphological trait | Seed |
| Heading date | Physiological trait | Flowering |
| Ligule length | Morphological trait | Culm leaf |
| Leaf length | Morphological trait | Culm leaf |
| Leaf width | Morphological trait | Culm leaf |
| Panicle length | Morphological trait | Dwarf |
| Seedling height | Morphological trait | Dwarf |

**Table 6:**
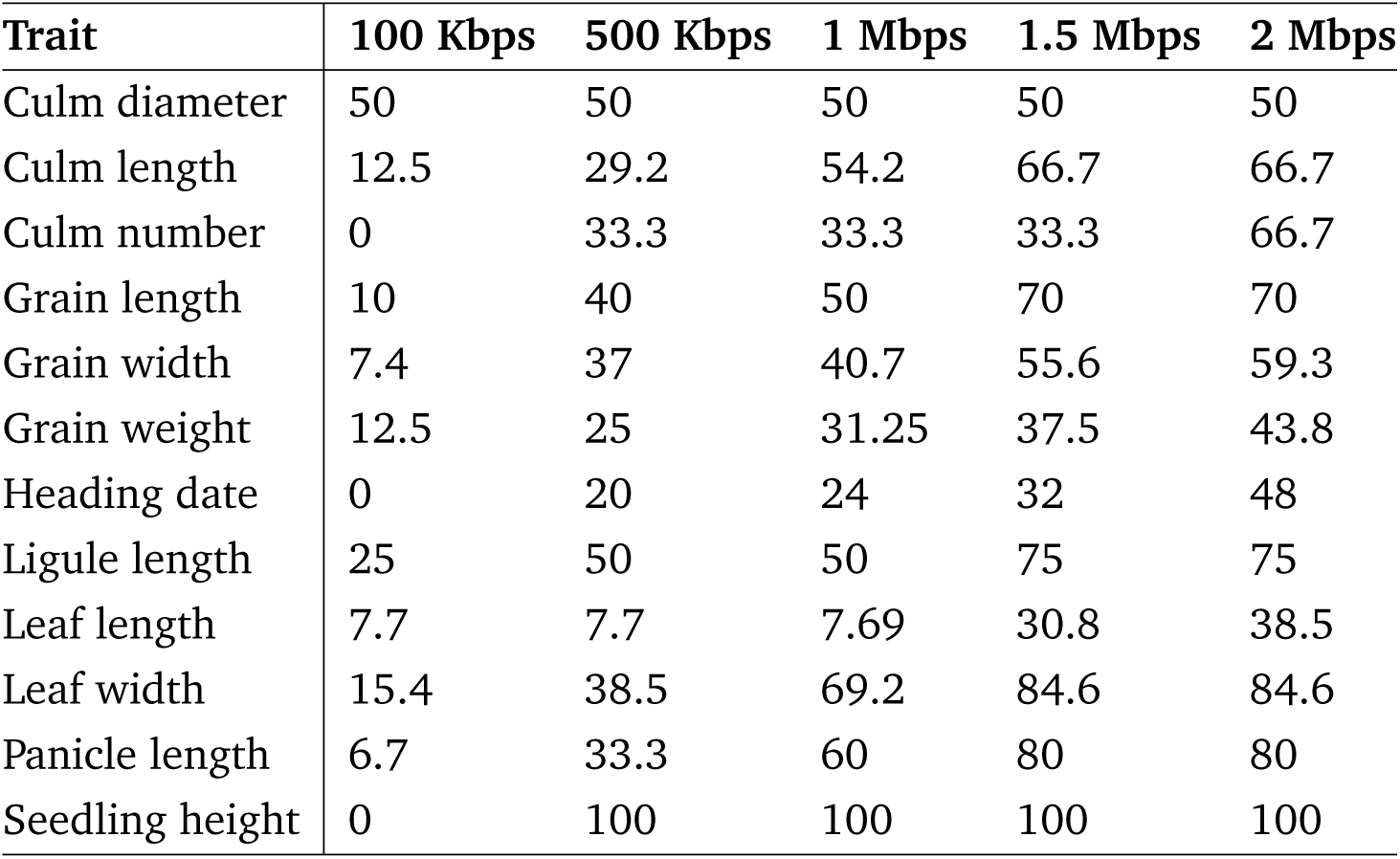
Percentage of selected markers (ICVM) in LD with a gene with major and minor characteristics in Table 5 at multiple distance thresholds. ICVM - Intersect cross validation markers.

| **Trait** | **100 Kbps** | **500 Kbps** | **1 Mbps** | **1.5 Mbps** | **2 Mbps** |
| --- | --- | --- | --- | --- | --- |
| Culm diameter | 50 | 50 | 50 | 50 | 50 |
| Culm length | 12.5 | 29.2 | 54.2 | 66.7 | 66.7 |
| Culm number | 0 | 33.3 | 33.3 | 33.3 | 66.7 |
| Grain length | 10 | 40 | 50 | 70 | 70 |
| Grain width | 7.4 | 37 | 40.7 | 55.6 | 59.3 |
| Grain weight | 12.5 | 25 | 31.25 | 37.5 | 43.8 |
| Heading date | 0 | 20 | 24 | 32 | 48 |
| Ligule length | 25 | 50 | 50 | 75 | 75 |
| Leaf length | 7.7 | 7.7 | 7.69 | 30.8 | 38.5 |
| Leaf width | 15.4 | 38.5 | 69.2 | 84.6 | 84.6 |
| Panicle length | 6.7 | 33.3 | 60 | 80 | 80 |
| Seedling height | 0 | 100 | 100 | 100 | 100 |

We observed that some of the selected associated markers in LD with known major controlling genes are at conservative distance thresholds (< 500 Kbps) while others are farther away (Table 7) for some of the traits considered. Examples of this are the grain size traits (length, width, and weight), and heading date. For the grain size traits, major genes such as *GS3* [38], *GS5* [24], *GW5* [43], *and GS6* [34] are in LD with an associated marker under the 500 Kbps threshold. However for heading date, although *Ghd7* [47] which is a major controlling gene is in LD with an associated marker under the 500 Kbps threshold, others such as *Hd6* [36, 48] are as far as ∼1.5 Mbps. We argue that being in LD with known controlling genes for the traits considered validates the selected associated markers. Furthermore, it also implies that not only is marker stability capable of validating known gene associations, but it can also be used in predicting novel associated genomic regions for a trait.

**Table 7:**
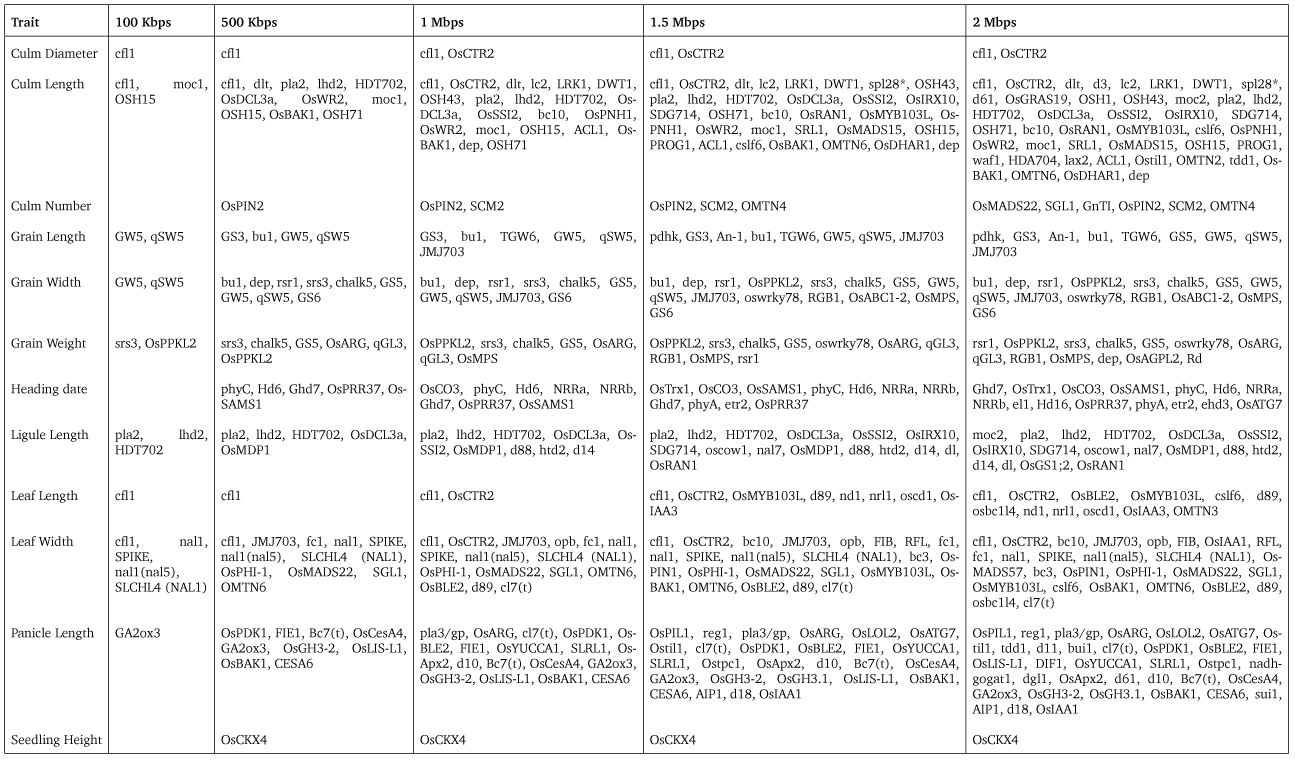
Genes with major and minor characteristics in Table 5 that are in LD with the ICVMs at different LD distance thresholds for each trait. ICVM - Intersect cross validation markers.

Co-functional network prediction of the annotated markers for grain size *(GS6, SGL*), heading (*phyC, OsSAMS1*) and culm length (*OsBAK1*) within the 500 Kbps distance threshold using RiceNet v2 [21], for example, also shows associations of the candidate markers to known genes, further corroborating the power of the proposed approach to identify regions that regulate target traits. For grain size, *GS6* (grain size 6, LOC Os06g03710) and *SGL* (short grain length, LOC Os05g06280) are connected to *JMJ703* (Jumonji C domain-containing protein 703, LOC Os05g10770; a histone H3K4-specific demethylase), *HDA703* (histone deacetylases 703, LOC Os02g12350) and *OsPPKL2* (protein phosphatase with Kelch-like repeat domain 2, LOC Os05g05240), all of which are involved in regulating seed development and morphology in rice. *GS6*, a unique member of the GRAS (which includes the first described-members GAI, RGA and SCR from which the name was derived) protein that encodes plant-specific family of transcription factors, is a negative regulator of grain size [34].

A study on *GS6* rice mutant showed that a pre-termination translation mutation in the *GS6* coding sequence resulted in an increase in grain width and weight [34]. *SGL*, a kinesin-like protein involved in the gibberellic acid biosynthesis pathway and response, regulates cell elongation and results in shorter grains and internodes in rice mutants [44]. Knockout studies on transgenic rice plants show pleiotropic effects of *JMJ703*, resulting in dwarf phenotype with reduced grain length, width and thickness [10] and repression of *HAD703*, also a member of histone deacetylases family, shortens rice peduncle and fertility [17]. *OsPPKL2* and *OsPPKL3* are the two homologs of *OsPPKL1*, which is encoded by *qGL3*. A transfer-DNA insertion mutant using *OsPPKL2* resulted in shorter grains, whereas *OsPPKL1* and *OsPPKL3* produced long grains [51].

For heading, gene network analysis clustered *phyC* with *phyB* and *OsEMF2b* (embryonic flower 2b), all of which are involved in photoperiod response, while *OsSAMS1* was grouped with *OsSAMS2, OsSAMS3* and *GF14c* (G-box factor 14-3-3c protein), and are known to regulate flowering time in rice. Functional characterization of the phytochrome gene family in rice, which only includes *phyA, phyB*, and *phyC*, has shown that mutation in either *phyB* or *phyC* alone or in double mutant combination of *phyA* with *phy B* or *phyC* resulted in early flowering under the long-day photoperiod [37] and that overexpression *OsEMF2b* resulted in early flowering by promoting the expression of key flowering genes in rice [45]. On the other hand, suppression of the SAMS (S-Adenosyl-l-methionine synthetase) genes, *OsSAMS1, OsSAMS2* and *OsSAMS3*, resulted in late flowering in rice while it is the overexpression of *GF14c* that showed delay in flowering time [23, 32].

For culm length, *OsBAK1* (a BRI1-associated receptor kinase) showed direct connection with brd2 (brassinosteroid-deficient dwarf2) and *Os4CL3*. Independent knockout studies on *brd2* and *Os4CL3* resulted in semi-dwarf phenotype and other defects in plant development [16] while overexpression of *OsBAK1* resulted in dwarf phenotype [22]. *OsBAK1* is the closest homolog of the Arabidopsis *BAK1* gene in rice and its overexpression enhances brassinosteroids signal and results in dwarf phenotype [42]. The results from gene network prediction based on direct neighborhood to known genes with similar function to those of candidate genes from our study further confirms the effectiveness of marker stability to identify true marker-trait associations in diverse rice landraces.

As further validation that the selected markers are informative we estimated the likelihood of randomly sampling markers in LD with the same genes as the selected markers. For 10,000 iterations we randomly sampled the same number of selected markers for each trait, checking if the genes in LD are the same as those selected using the proposed procedure. We estimated the likelihood by dividing the number of times the randomly sampled markers are in LD with the same genes as the selected markers by the total number of iterations. We performed this procedure for the five thresholds considered. The assumed null hypothesis was that the same, or greater, number of genes in LD with the selected markers would be found by randomly sampled markers. With a significance level of 0.01, the results reject the null hypothesis (Table 2).

Given the extensive validation procedures performed on the selected associated markers, we argue that these results suggests that the selected markers that are not in LD known controlling QTLs reveal previously unknown interacting regions which control the traits considered, and serves as novel knowledge that can be integrated into future rice breeding efforts. Furthermore, we argue that these results are particularly important given the diversity of the samples in the population study, as the selected associated markers are significant irrespective of sample variety.

### Comparison to GEMMA

On the twelve agronomic traits a GEMMA linear mixed model was performed with kinship matrix and the first three principal components as covariates. Both kinship and the principal components were computed from the same subset of SNPs and all 3023 varieties present in the dataset were used. Multiple testing correction was performed using false discovery rate. At a significance level of −*log*_10_(1*e* − 5), GEMMA identified a total of eleven significant SNPs for three of the twelve traits considered; six for grain length, four for grain width and a single SNP for grain weight (Fig. 2). These SNPs are in LD with only *GS3* and *GW5* at 500 Kbps. It is worth noting that the proposed procedure also selects two of the six SNPs identified for grain length, all four SNPs identified for grain width but does not select the single SNP identified for grain weight. However, rather than replacing standard approaches like GEMMA with marker stability, we argue that they should be used in conjunction, with one method serving as validation for other and identifying associated markers that the other might miss.

**Figure 2:**
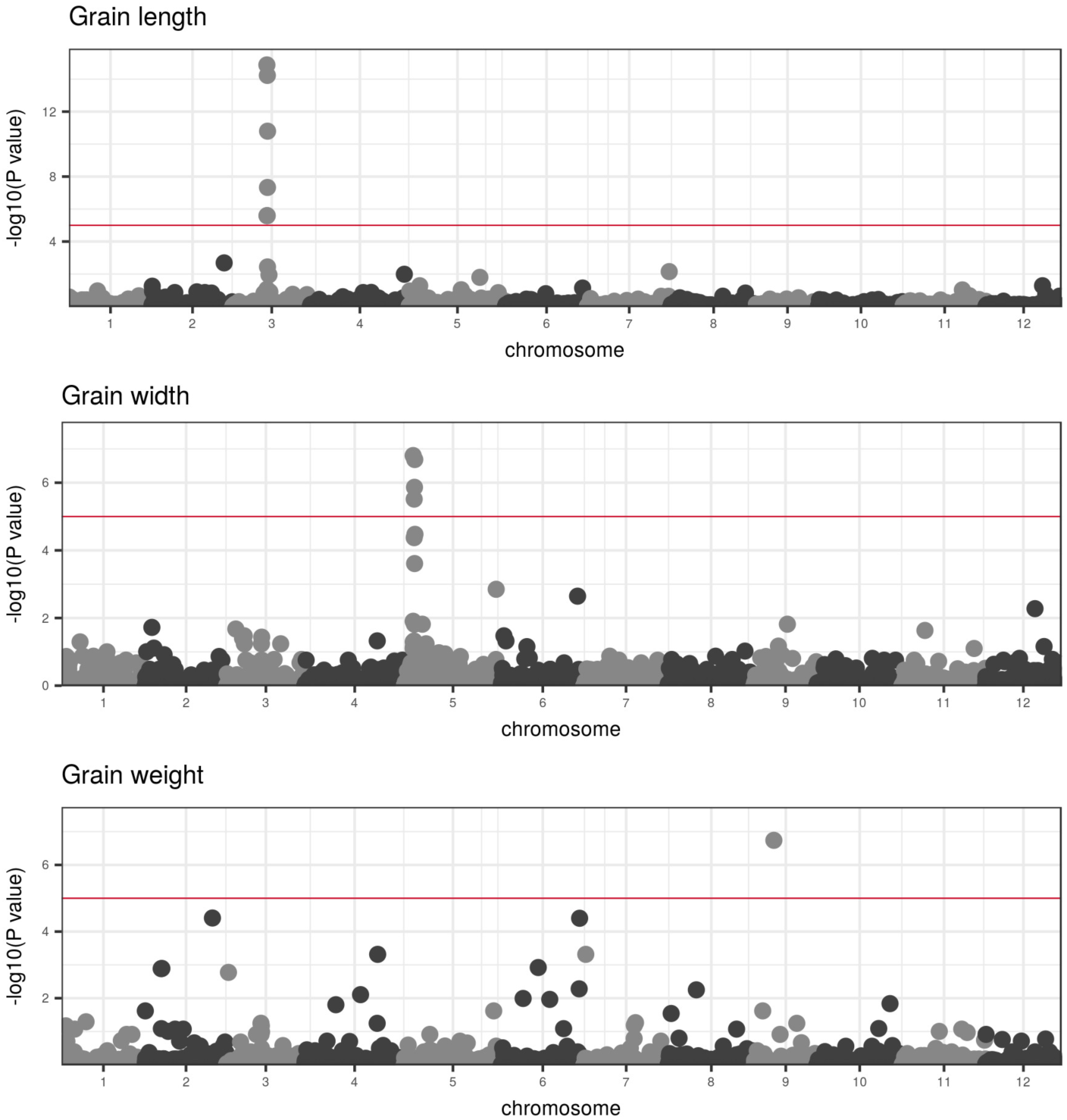
Manhattan plots for grain length, grain width and grain weight after false discovery rate correction at a signficance level of −*log*_10_(1*e* − 5).

We also compared our approach to GEMMA on a hundred simulated traits. The GEMMA SNPs were ranked by p-values and we ranked the ICVM markers using RF. We then selected the ten most significant SNPs identified by both methods and computed Sum10 – the sum of their effects taken from the original model behind the traits. This reflects the realistic scenario where around ten SNPs can be selected for experimental validation. For some traits, marker stability selected fewer than ten SNPs, in such cases we used the same number of SNPs for GEMMA. See Fig. 3(A) for the distribution of the number of markers selected by marker stability. Sum10 analysis showed that GEMMA had a higher Sum10 value for 21 of the simulated traits, GEMMA and ICVM had the same value for 2 traits and ICVM had a higher value for 77 traits (Fig. 3(B) and Fig. 3(C)). The assumed null hypothesis was that there is no difference in performance between the two methods, paired 2-tail t-test resulted in a p-value of 2.037*e* − 07 (t-value = 5.587, degrees of freedom = 99). Therefore the null hypothesis can be rejected with a significance level of 0.01.

**Figure 3:**
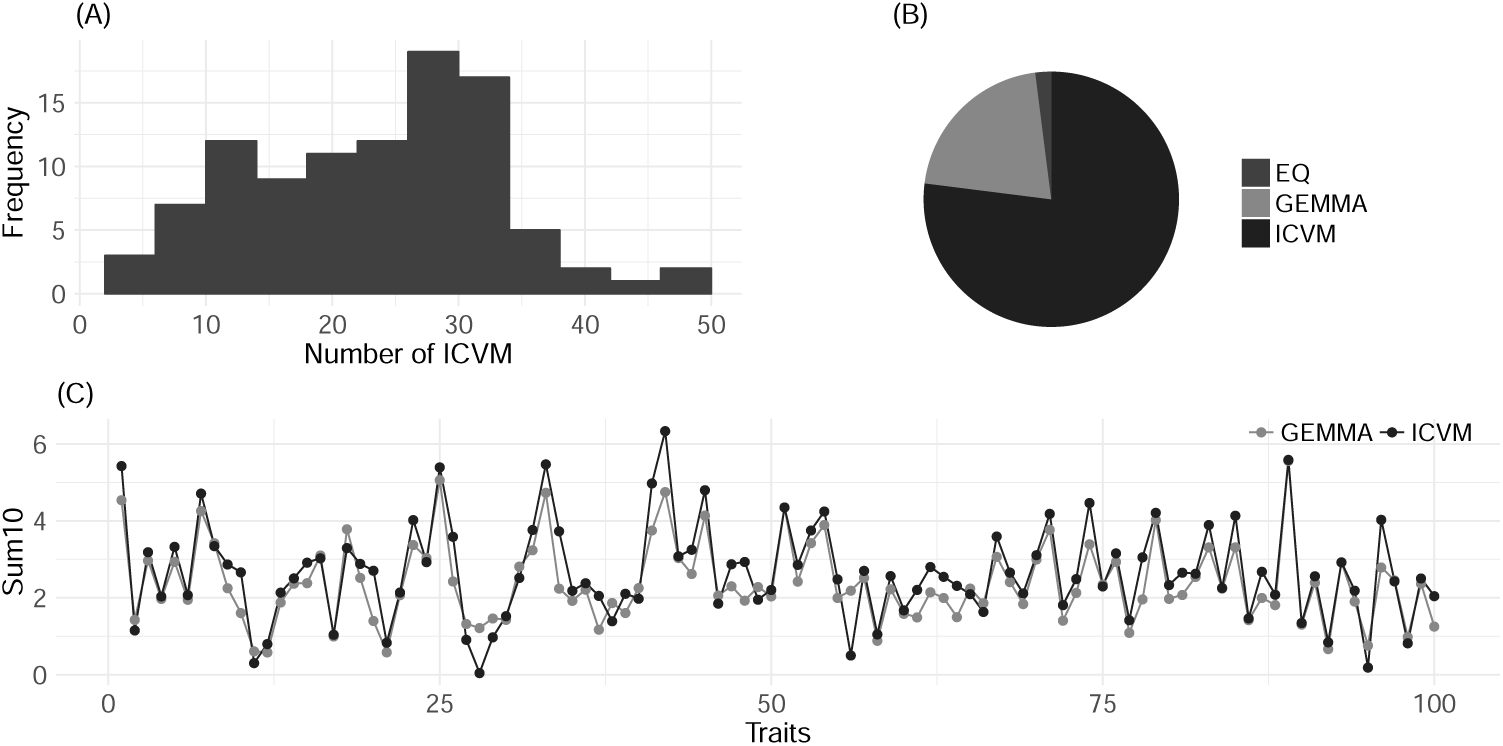
Graphical representations of the results from the simulation experiments. **(A)** shows the distribution of the number of markers selected using the proposed approach. **(B)** shows the proportion of the simulated traits for which GEMMA and ICVM performed best on Sum10 analysis. Eq is the proportion for which GEMMA and ICVM performed equally. **(C)** shows the Sum10 values for GEMMA and ICVM on the simulated traits.

**Figure 4:**
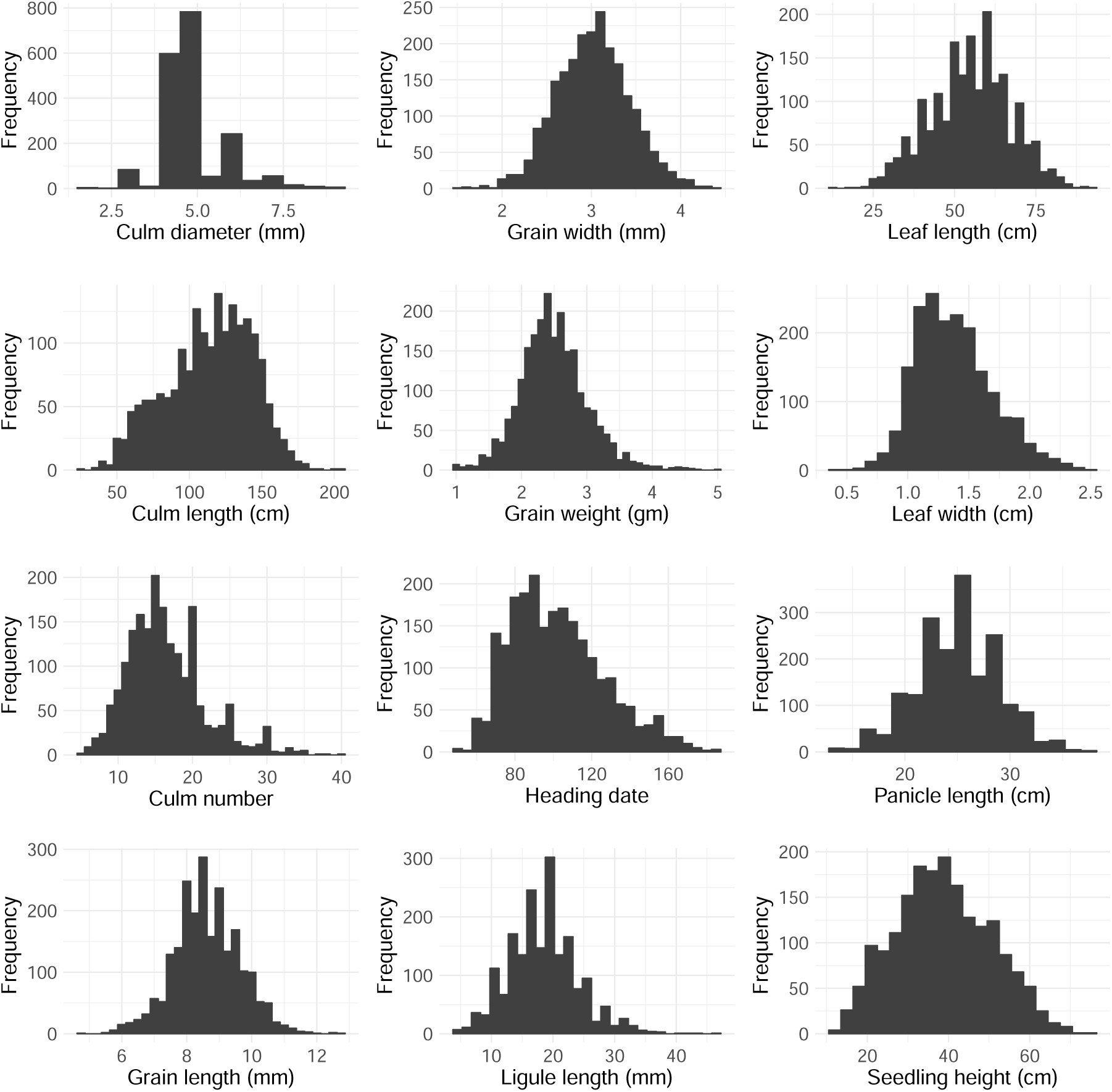
Distribution of traits used in this study.

## Conclusion

We have reported our findings from performing trait prediction and association mapping by marker stability using LASSO. Our results show that marker stability can be used to identify associated markers in diverse rice populations which can be used for trait prediction and association mapping. For trait prediction, we found that one might need to make a trade-off between model size and predictive power. For association mapping, we found that marker stability typically outperforms traditional mixed-model methods. Therefore, we conclude that marker stability should be used in conjunction with traditional mixed-model analysis for the identification of associated markers in rice breeding efforts with diverse populations, which could improve our knowledge of the genetic underpinnings of complex agronomic traits.

